# Conserved and tissue-specific immune responses to biologic scaffold implantation

**DOI:** 10.1101/2023.08.15.553390

**Authors:** Sabrina DeStefano, Aditya Josyula, Mondreakest Faust, Daphna Fertil, Ravi Lokwani, Tran B. Ngo, Kaitlyn Sadtler

## Abstract

Upon implantation into a patient, any biomaterial induces a cascade of immune responses that influences the outcome of that device. This cascade depends upon several factors, including the composition of the material itself and the location in which the material is implanted. There is still significant uncertainty around the role of different tissue microenvironments in the immune response to biomaterials and how that may alter downstream scaffold remodeling and integration. In this study, we present a study evaluating the immune response to decellularized extracellular matrix materials within the intraperitoneal cavity, the subcutaneous space, and in a traumatic skeletal muscle injury microenvironment. All different locations induced robust cellular recruitment, specifically of macrophages and eosinophils. The latter was most prominent in the subcutaneous space. Intraperitoneal implants uniquely recruited B cells that may alter downstream reactivity as adaptive immunity has been strongly implicated in the outcome of scaffold remodeling. These data suggest that the location of tissue implants should be taken together with the composition of the material itself when designing devices for downline therapeutics.

## 2. INTRODUCTION

The immune response to biomaterial scaffolds has been long appreciated as a critical mediator in scaffold remodeling and integration with surrounding tissues. Various immune cells have been implicated in these responses, including macrophages that help degrade collagenous matrices (both exogenous and natural tissue) and remodel tissue-associated extracellular matrix in trauma and development [1]. Macrophages have been observed in the tissue development of regenerative organisms such as axolotls, which depend on macrophages for limb regeneration [2]. In addition to macrophages, eosinophils have been implicated in muscle and liver regeneration [3, 4]. Adaptive immune cells such as T cells, specifically Th2-polarized CD4^+^ T cells, have been associated with positive outcomes in wound healing, macrophage polarization, and subsequent tissue remodeling in a murine model of volumetric muscle loss [5]. Type-2 immune responses correlate with myotube formation and prevent excessive fibrosis and adipogenesis [3, 6]. This has been shown specifically with skeletal muscle tissue with the cytokine interleukin 4 (IL-4) that acts as both a myoblast recruitment factor and induces fusion to form the multinucleate muscle fibers through NFATC2 [6]. In contrast, pro-fibrotic materials such as polyethylene induce a more type-1 biased inflammatory microenvironment that promotes neutrophilic inflammation followed by dense collagen deposition and fibrosis [7]. This follows the canonical foreign body response, characterized by protein adsorption to the material followed by macrophage infiltration and an attempt at degradation and phagocytosis, which ultimately results in fibrotic capsule formation when the material cannot be degraded [8, 9]. The immune system mediates these pro-healing and pro-fibrotic responses playing a critical role in the positive and pathogenic outcomes of implanted materials [10, 11].

Biologic scaffolds are used clinically in various tissue locations for application in the presence and absence of injured tissue. Decellularized extracellular matrix (ECM) scaffolds are used for abdominal wall repair during hernia reconstruction, dural repair after neurosurgery, breast filling after lumpectomy, diabetic foot ulcer treatment, skin injury reconstruction, and have been tested for the treatment of volumetric muscle loss (VML) and more significant tissue defects [12, 13]. These contexts come with different immune microenvironments ranging from immune-privileged sites (such as the brain in dural repair) to abutting visceral organs (in abdominal wall repair) to barrier tissues (in skin wound repair). Specific tissue locations, such as the skin, can withstand strong inflammatory responses with dense scar tissue deposition without the risk of loss of life. However, other tissue sites, such as the abdominal cavity, can induce fibrotic responses that lead to diseases associated with organ failure and death, like liver cirrhosis, intestinal fibrosis, pancreatic fibrosis, and others [14]. Different tissue sites have developed diverged propensities for immune polarization and activation to suit the location where an immune challenge occurs [15]. These differences are due to both the surrounding stromal and parenchymal cells, as well as the profiles of resident immune cells. Tissue-resident macrophages vary greatly depending on tissue location. For example, microglia (brain-resident macrophages) differ from liver-resident macrophages (Kupffer cells) and skin-resident cells. These cells have different epigenetic profiles and propensities for immune polarization [16].

Tissues have previously been viewed as passive recipients of immune protection, recent work has appreciated the active role that stromal and parenchymal cells play in influencing and generating immune activity [15]. Many cell types can secrete immune-active proteins and chemicals that alter the immune activation and polarization in response to a given stimulus. Cell-cell communications, such as macrophage-fibroblast crosstalk, are critical in the foreign body response [17]. In the context of biomaterial implantation, it has been previously reported that the location of a hydrogel implant can alter its host response as determined by histologic examination [18]. These factors suggest that tissue context is important in biomaterial responses and outcomes. While there are several mouse models available for different biomaterial applications, many studies use standardized *in vitro* and subcutaneous implantation models to evaluate biocompatibility. There is a large gap in knowledge between preclinical studies and clinical implementation due to variables such as tissue implant location that are not always considered. Therefore, this study evaluates the immune response to a biomaterial in different body locations (intraperitoneal versus subcutaneous) and in the presence or absence of an injury (subcutaneous non-traumatic versus subcutaneous traumatic muscle injury). The findings from this study provide a better understanding of the implant site’s immune environment, which will help design biomaterials for more diverse clinical applications.

## 3. MATERIALS AND METHODS

### 3.1 Decellularized extracellular matrix synthesis

Decellularized extracellular matrix (ECM) was synthesized as previously described. Briefly, the small intestine of Yorkshire Pigs was isolated, and the mucosa and muscular layers were physically removed from the submucosa connective tissue layer (SIS). The resulting submucosa layer was rinsed in distilled water and then incubated in antibiotic (PenStrep, Gibco) diluted in distilled water at 4°C overnight to remove residual mucosal debris. SIS was rinsed thoroughly in distilled water and then transferred to a sterile container with 0.1 % peracetic acid (Sigma) and 4% ethanol (Sigma) diluted in sterile distilled water and incubated with vigorous stirring for 30 minutes. The resulting ECM was rinsed in successive washes of sterile water followed by sterile 1xPBS until the tissue was neutralized. Decellularization was confirmed with dsDNA quantification and histologic evaluation. ECM was rinsed with a final sterile distilled water wash, drained of liquid, and frozen at -80 °C before lyophilization and cryogenic milling to form a powder. The powder was hydrated in sterile surgical saline on the day of surgery and loaded into 1 mL syringes.

### 3.2 Mouse models of biomaterial implantation

Six (6) to 8-week-old wild-type female C57BL/6 mice were sourced from Jackson Laboratories. After 1 week of equilibration in the facility, animals for volumetric muscle loss (VML) were anesthetized under 2.0% isoflurane, and hair was removed from hindlimbs with an electric razor followed by depilatory cream. The following day, mice for all groups were anesthetized and implanted with materials. All following procedures were conducted under an approved animal protocol reviewed by the NIH Clinical Center Animal Care and Use Committee in compliance with the NIH Guide for Care and Use of Laboratory Animals.

#### 3.2.1 Intraperitoneal (IP) implants

after wiping the ventral area with a 70% isopropanol-soaked gauze pad, an 18 G needle was inserted along the linea alba and 50 μL (for FACS studies) or 200 μL (for histologic studies) of ECM was injected.

#### 3.2.2 Subcutaneous (SQ) implants

after the dorsal surface of the mouse was wiped down with 70% isopropanol, two 50 μL implants were created under the skin along the spine.

#### 3.2.3 Volumetric muscle loss (VML) implant

after sterilizing with 3 successive rounds of betadine followed by isopropanol, a 1 cm incision was made in the skin overlying the quadriceps muscle group, and the fascia was dissected away to reveal the muscle. A 3 mm portion of the muscle (corresponding with 30 mg of tissue) was excised, and the resulting defect was filled with 50 μl of ECM. The skin was closed using 7 mm wound clips, and the procedure was repeated on the contralateral leg.

### 3.3 Flow cytometry

After 7 and 21 days after implantation, animals were euthanized with carbon dioxide. For SQ implants, the skin was excised around the implant area, and the implant with the associated capsule was examined away from the dermis. For VML implants, the quadriceps with ECM material was dissected out along the femur to the hip. The skin and abdominal wall were dissected carefully for IP implants to prevent bleeding. The IP cavity was lavaged with 1 mL of serum-free RPMI, and then any visible ECM was removed from the IP cavity. The resulting tissue isolates were digested in 0.25 mg/ml Liberase TM with 0.2 mg/ml DNase I in serum-free RPMI for 45 minutes at 37°C on a shaker at 100 rpm at a total volume of 5 mL per 50 μl implant. Digested samples were filtered through a 70 μm cell strainer and rinsed with room temperature 1xPBS. Samples were spun down at 350xg at room temperature for 5 minutes, and the resulting pellet was washed twice in 1xPBS before staining for 30 minutes in LIVE/DEAD Blue viability dye or 7-AAD. As previously described, the cell pellet was washed three times in cold 1xPBS before staining in a surface antibody cocktail (**Supplemental Tables 1 and 2**,[19-21]). Samples were washed three times and then run on a Cytek Aurora 5 laser spectral flow cytometer. Single color controls for unmixing were made using peripheral blood mononuclear cells.

### 3.4 Histology

IP implants were kept intact within the peritoneum. The overlying skin was removed to isolate the entire peritoneal cavity, fixed for 48 hours in Bouin’s Solution, then dissected into three 1 cm sections before being fixed for another 24 hours. Bouin’s solution was rinsed out with successive rinses of 1xPBS before being placed in 70% ethanol for FFPE processing.

SQ implants were dissected from the underlying muscle with overlying skin attached to the implant. Samples were fixed overnight in 10% neutral buffered formalin (NBF) overnight, and then rinsed in distilled water before being placed in 70% ethanol for formalin-fixed paraffin-embedded (FFPE) processing.

VML implants were dissected and collected similarly to those previously described for flow cytometry. Samples were incubated in 10% NBF for 48 hours before rinsing in distilled water and placing in 70% ethanol for FFPE processing. All models were dehydrated in a graded ethanol series and cleared in xylenes before paraffin infiltration using an automated tissue processor (Leica). Samples were mounted on paraffin blocks, and 5 μm sections were taken before being baked for 3 hours at 60°C and stained with hematoxylin and eosin (H&E, Sigma) or Masson’s trichrome (Sigma) as per manufacturer’s instructions. Slides were imaged on an EVOS microscope.

### 3.5 Data Analysis

Flow cytometry data was unmixed on SpectroFlo using single color controls and then exported as .fcs files for analysis on FlowJo v 10.9.0. Gates were set using fluorescence-minus-one controls. Data were exported from FlowJo, and statistical analysis was completed on GraphPad Prism v 9.5.1. tSNE (t-stochastic neighbor embedding) clustering was performed via FlowJo [22]. Histology images of the implants from H&E-stained slides were opened on FIJI (ImageJ v 2.9.0), split channels to isolate the hematoxylin stain, then converted to 16-bit greyscale, and thresholded to isolate nuclei. Nuclei were counted using the Analyze Particles plugin, and events were counted if they were above 50 pixels in the area to remove the artifact. The resulting data were analyzed in GraphPad Prism v 9.5.1.

## 4. RESULTS

### 4.1 Macrophage and eosinophil infiltration dominates the myeloid response to the ECM material regardless of implant locations

To evaluate the role of tissue location in immune responses to naturally-derived biomaterial scaffolds, we implanted 50 μl of a particulate decellularized extracellular matrix (ECM) in the intraperitoneal space (IP), the subcutaneous space (SQ), and a volumetric muscle loss skeletal muscle injury (VML). The resulting scaffolds were excised and processed for immunologic analyses via a 21-color immunophenotyping panel focused on the innate immune response to materials (**Fig. 1a, Supplemental Figure 1**). We were able to identify a variety of different immune cell types in response to biomaterials in various tissue locations, including macrophages (F4/80 and/or CD68^+^), neutrophils (Ly6G^+^), eosinophils (Siglec-F^+^), basophils (CD200R3^+^), type 1 conventional dendritic cells (CD103^+^XCR1^+^MHCII^+^), and other non-myeloid antigen-presenting cells (APCs, Lin^-^MHCII^+^) (**Fig. 1b**). To analyze the data beyond the standard population identifications applied with manual gating, we used dimensionality reduction algorithms including t stochastic neighbor embedding (tSNE) to visualize heterogeneity of myeloid cell populations (**Fig. 1c**). In addition to confirming the manually identified myeloid cell populations, subpopulations of macrophages, eosinophils, basophils, monocytes, and antigen-presenting cells (MHCII+) were identified on islands associated with different tissue locations.

**FIG 1.**
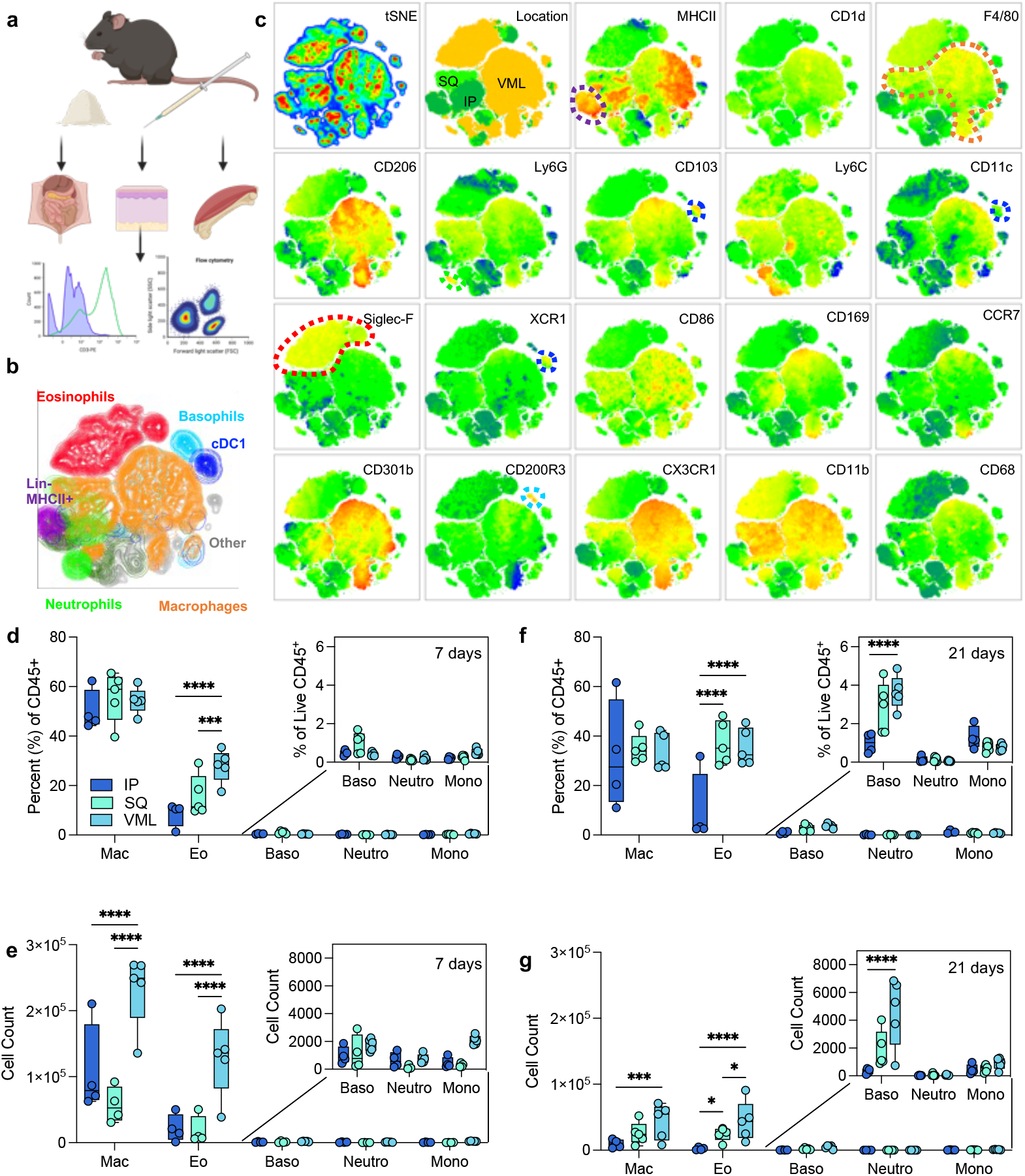
Myeloid response to biologic scaffolds in different tissue locations. (a) Experimental workflow (b) tSNE of manually gated immune cell (CD45+) populations in ECM scaffolds (c) Expression of myeloid phenotyping markers in different islands. SQ = subcutaneous implant; IP = intraperitoneal implant; VML = volumetric muscle loss skeletal muscle injury implant. (d) Immune cell populations as a percent of live immune cells at 7 days post-implantation (dpi). (e) Immune cell counts at 7 dpi from 50 μl implants. (f) Immune cell populations as a percent of live immune cells at 21 dpi. (g) Immune cell counts at 21 dpi. Data are range, n = 4 – 5, ANOVA with Tukey posthoc. * = p < 0.05; *** = p < 0.001; **** = p < 0.0001. Schematic made with BioRender.

**FIG 2.**
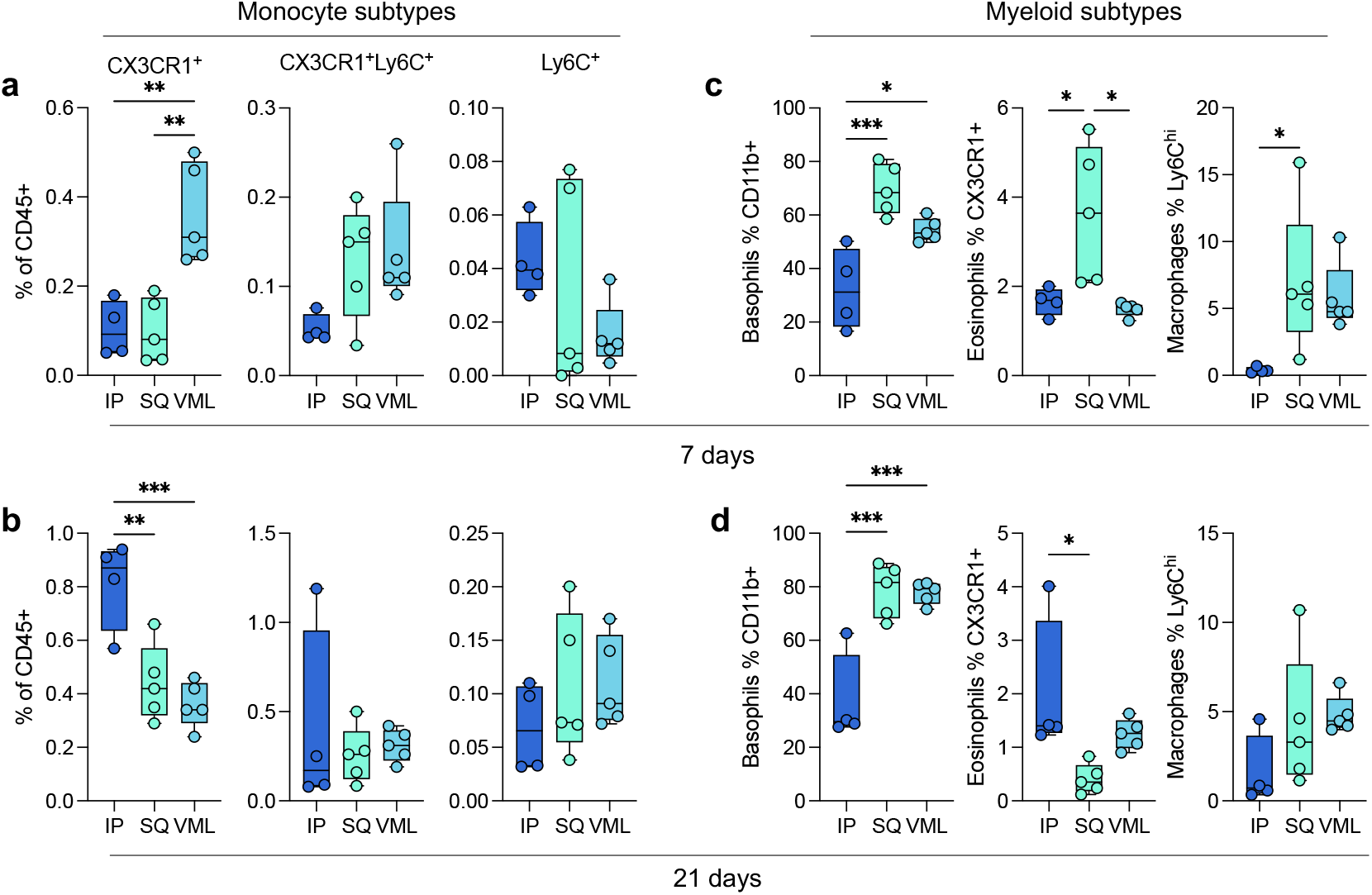
Myeloid subtypes vary in an implant location and time-dependent manner. (a) Monocyte subtypes at 7 dpi. (b) Monocyte subtypes at 21 dpi. (c-d) Basophil, eosinophil, and macrophage subtypes at (c) 7 dpi and (d) 21 dpi. Data are range, n = 4 – 5, ANOVA with Tukey posthoc. * = p < 0.05; ** = p < 0.01; *** = p < 0.001

At 7 days post-injury, all implants comprised macrophages as 50% of total immune infiltrate 50.15% ± 4.12% IP; 55.94% ± 4.54% SQ; 54.3% ± 2.36% VML). There were a significantly higher proportion of eosinophils as a fraction of immune infiltrate in subcutaneous space with and without tissue injury in comparison to intraperitoneal implantation (**Fig. 1d**). This held to 21 days post-implantation, wherein there was an increased fraction of eosinophils when compared to macrophages which decreased between 7- and 21-days post-implantation. There was also a modest increase in basophils at 21 days post-implantation in the subcutaneous and VML models, which was significantly higher than IP implantation (**Fig. 1f**). In terms of the count of cells per implant, the muscle injury significantly increased the number of macrophages and eosinophils that responded to the biomaterial scaffold which persisted out to 21 days (**Fig. 1e-g**).

### 4.2 Multiple myeloid subtypes vary in tissue location-dependent manner

In addition to the main myeloid cell types, several subtypes were identified. As previously mentioned, a number of islands were detected in the tSNE visualization of flow cytometry data suggesting the presence of sub-populations of multiple immune cells. Early in response to biomaterials, there was significant recruitment of CX3CR1^+^ monocytes in the VML microenvironment (0.36% ± 0.05%), as opposed to the IP location that had higher CX3CR1^+^ monocytes at 3 weeks post-implantation (0.81% ± 0.08%; **Fig. 1a**,**b**). These monocytes have been associated with a pre-M2 phenotype, and M2-like macrophages have been associated with biological scaffold remodeling. In addition to different monocytic infiltration, we saw increases in CD11b^+^ Basophils at 7 and 21 days post-implantation in both SQ and VML models compared to **Fig. 1c**,**d;** 7 days 32.23% ± 7.58% IP, 69.6% ± 4.22% SQ, 54.44% ± 1.92% VML; 21 days 37.33% ± 8.44% IP, 78.58% ± 4.43% SQ, 77.74% ± 1.84% VML; *p* < 0.05). These cells may also be mast cells which are more tissue-resident but also express the CD200R3 marker. Both basophils and mast cells play a role in type-2 polarized immune responses, specifically allergy and asthma. We saw an early preference for CX3CR1^+^ eosinophils in SQ implants, which was highest in IP implants at 21 days post-implantation (2.01% ± 0.67%). We also saw a higher proportion of Ly6C^hi^ macrophages (CD11b^+^ and CD68 or F4/80^+^) in subcutaneous implants at 7 days post-implantation compared to IP implants (7.02% ± 2.42% v 0.39% ± 0.11%, *p* = 0.0469).

### 4.3 Injury induces a strong M2-like polarization in response to ECM implants

Though macrophage polarization does not follow a binary nor fit into easily categorizable phenotypes, we sought to understand the macrophage activation profiles and what possible function of macrophages at the different tissue locations. In this study, we used the expression of two M1-associated markers (CD86, a co-stimulatory molecule, and CCR7, a chemokine receptor mediating lymph node recruitment) and two M2-associated markers (CD206, the mannose receptor, and CD301b, a scavenger receptor associated with phagocytosis of Gal-GalNAc-modified antigens) (**Fig. 3**). The latter (M2 macrophages) have previously been associated with positive outcomes in biologic scaffold remodeling. These markers were evaluated for several myeloid cell populations, including Ly6C^hi^ macrophages, Ly6C^lo^ MHCII^+^ macrophages, Ly6C^lo^ MHCII^-^ macrophages, and cDC1s. The strongest differences in expression were seen with CD206 and CD301b expression on MHCII-macrophages (**Fig. 3a;** 2.84 x MHCII+ macrophages, 7d VML, *p* < 0.0001). This expression was greater at 21 days post-injury than 7 days post-injury. These cells had a lower expression of CD86 than their MHCII+ counterparts (0.46-fold, 21d VML) suggesting the latter are the main antigen-presenting macrophages. cDC1s also expressed high levels of CD86 compared to other markers supporting their role as canonical APCs. Injury induced a more robust CD86 expression on MHCII^+^ macrophages at 7 and 21 days post-implantation, whereas cDC1s expressed the highest CD86 expression in IP implants (**Fig. 3b**,**c**). MHCII^-^ macrophages expressed the highest CCR7 in VML contexts at both time points (**Fig. 3d**,**e**). Regarding M2-associated markers, VML implants induced the highest expression of CD206 (**Fig. 3f**,**g**) and CD301b (**Fig. 3h**,**i**) compared to the non-trauma applications in IP and SQ applications.

**FIG 3.**
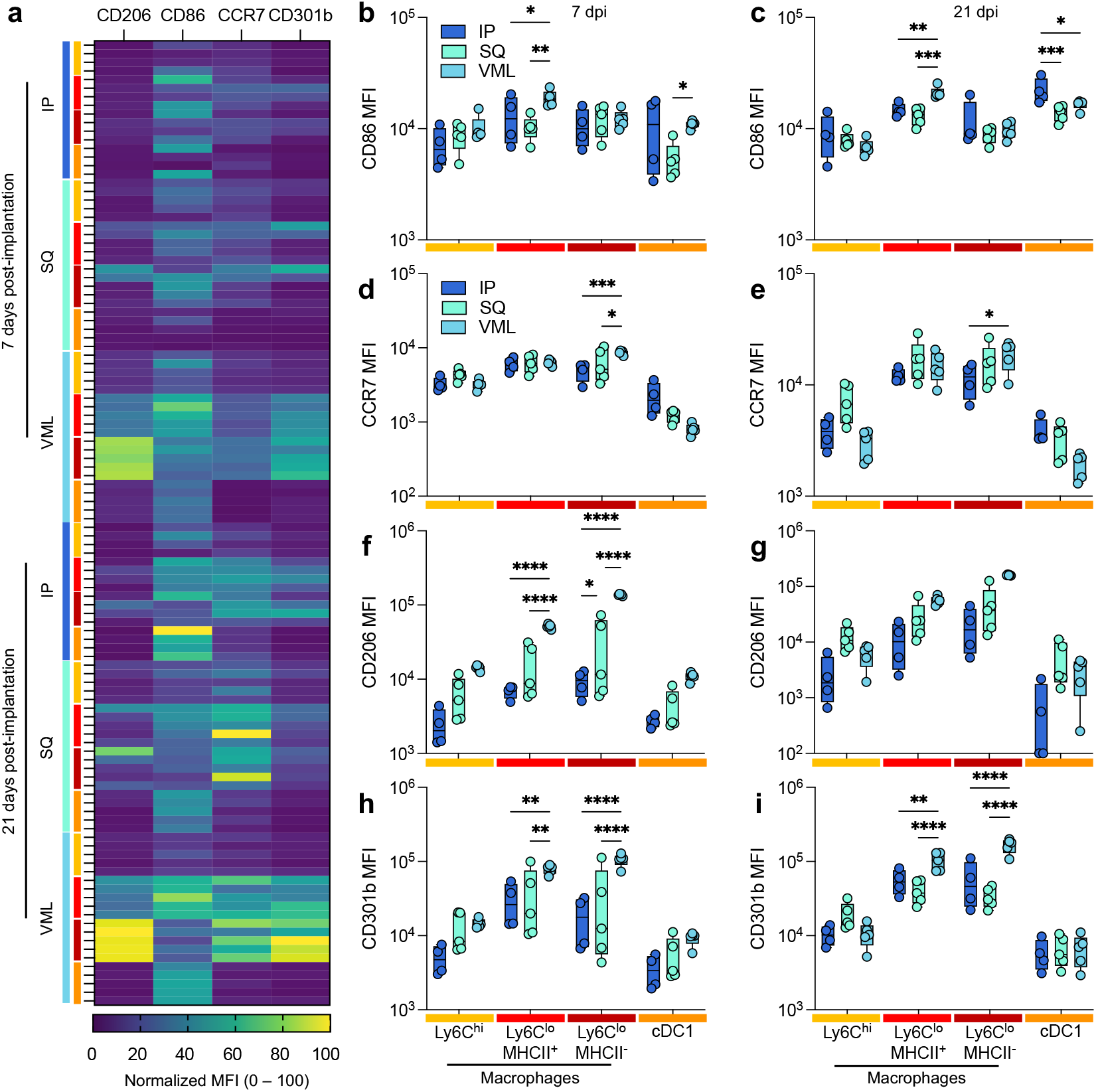
Injury induces a stronger type-2 polarized immune response than non-traumatic applications. (a) Myeloid polarization markers CD206, CD301b (M2-like), CD86, CCR7 (M1-like) over time. (b-c) CD86 median fluorescence intensity (MFI) at (b) 7 days post-implantation (dpi) and (c) 21 dpi. (d-e) CCR7 MFI at (d) 7 dpi, and (e) 21 dpi. (f-g) CD206 MFI at (f) 7 dpi and (g) 21 dpi. (h-i) CD206 MFI at (h) 7 dpi, and (i) 21 dpi. Yellow = Ly6C^hi^ macrophages, Red = Ly6C^lo^ MHCII^+^ Macrophages, Dark Red = Ly6C^lo^ MHCII^-^ macrophages, Orange = type 1 conventional dendritic cells (cDC1s). Data are range, n = 4 – 5, ANOVA with Tukey posthoc. * = p < 0.05; ** = p < 0.01; *** = p < 0.001; **** = p < 0.0001.

### 4.4 Antigen-presenting cells are strongly dependent on tissue location

Various antigen-presenting cells are present in response to ECM scaffolds. These include macrophages, dendritic cells, and a lineage negative MHCII^+^ population (**Fig. 4**). At 7 days post-injury Ly6C^hi^ macrophages had the highest MHCII^+^ population in SQ implants compared to IP and VML tissue locations. Ly6C^lo^ macrophages still had a significant MHCII^+^ population, with all macrophage subtypes having 20 – 60% of cells positive for the antigen presentation complex (**Fig. 4a**). All tissue locations recruited cDC1s, as determined by the expression of MHCII, CD103, XCR1, and CD11c (**Fig. 4b**). An unidentified negative lineage (CD11b^-^CD11c^-^Ly6C^-^CD68^-^F4/80^-^ SiglecF^-^CD200R3^-^Ly6G^-^) with MHCII^+^ cell population with a lymphocyte-like scatter profile was robustly upregulated in response to IP implants but absent in response to SQ and VML tissue locations (**Fig. 4c**).

**FIG 4.**
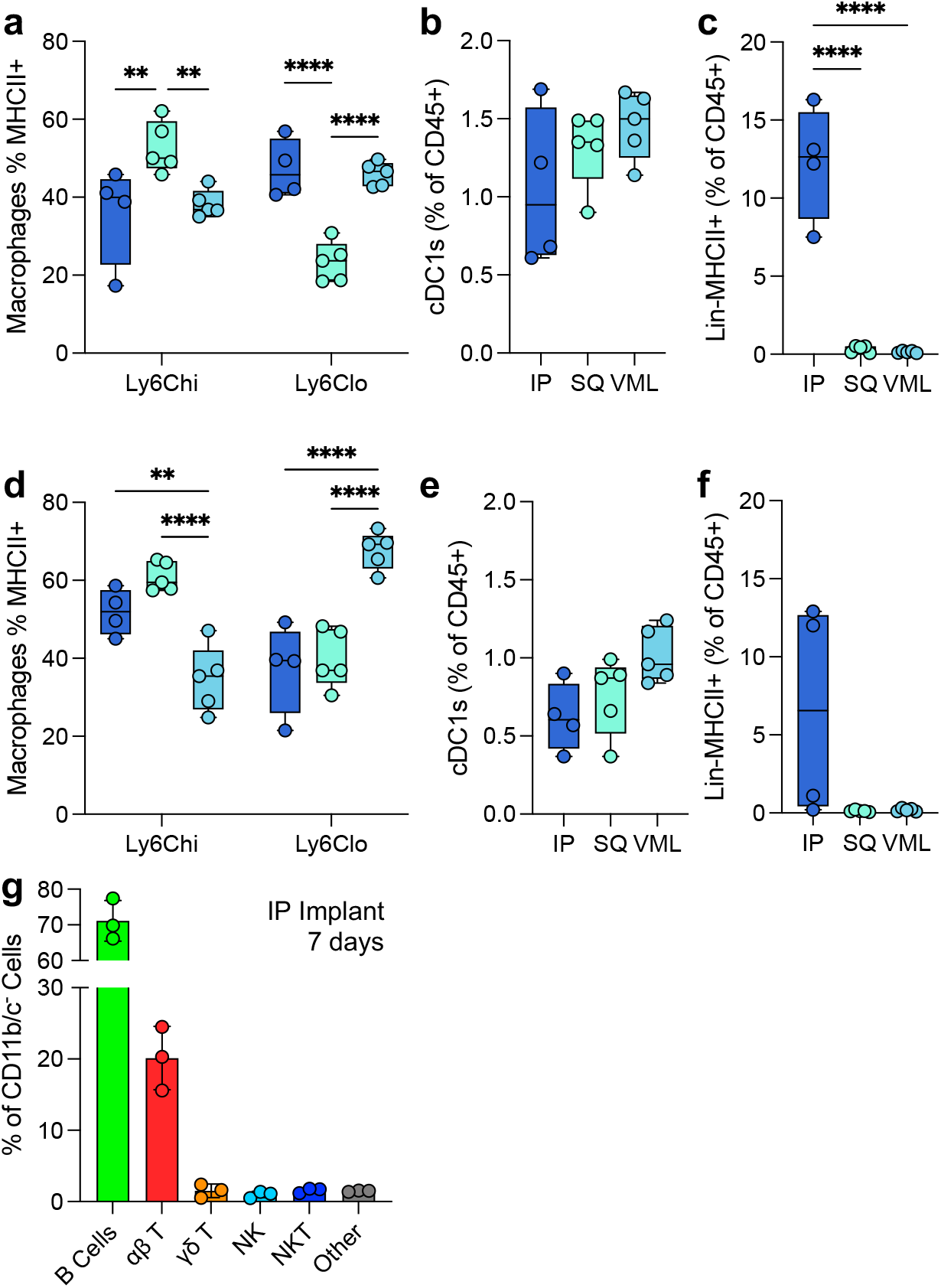
Variability in myeloid and lymphoid antigen-presenting cells in the tissue microenvironment. (a) The proportion of macrophages expressing MHCII at 7 dpi. (b) The proportion of total immune cells that are cross-presenting cDC1s at 7dpi (c) The proportion of lymphoid-like Lin-MHCII+ unidentified antigen-presenting cells 7dpi. (d-f) The proportion of (d) MHCII^+^ macrophages, (e) cDC1s, and (f) Lin^-^MHCII^+^ cells at 21dpi. (g) Lymphoid profile of IP implants. Data are range (a-f) or mean ± standard deviation, n = 3 – 5, ANOVA with Tukey posthoc. ** = p < 0.01; *** = p < 0.001; **** = p < 0.0001.

These patterns persisted to 21 days post implantation with increases in MHCII expression on Ly6C^hi^ macrophages for IP and SQ implants and increases in MHCII expression on Ly6C^lo^ macrophages in VML implants (**Fig. 4d**). cDC1s persisted but decreased by 21 days post-implantation, as seen previously in VML implants (**Fig. 4e**; 0.59-fold IP, 0.58-fold SQ, 0.70-fold VML). The lineage-negative cell population was still present in some mice tested at 21 days post-injury (**Fig. 4f**). To identify this Lin^-^MHCII^+^ population, we evaluated the lymphocytic profile of IP implants with a flow cytometry panel quantifying the presence of B cells (B220^+^), αβ T cells (TCRβ^+^NK1.1^-^), NK cells (NK1.1^+^), NKT cells (TCRβ^+^NK1.1^+^), and γδ T Cells (TCRβ^-^TCRγδ^+^, **Supplemental Figure 2**). Here, we found a robust preference for B cell recruitment (70% of total CD11b^-^CD11c^-^ cells) followed by αβ T cells (20%; **Fig. 4g**). Whereas in our previously published works, we found that the main lymphocytic cells responded to ECM implants in VML are αβ T cells and NK cells with very few B cells [19].

### 4.5 Cellular density within the material implant is dependent upon tissue location

Histologically, the structure and cellular infiltration of the materials varied greatly depending upon the tissue location (**Fig. 5**). Intraperitoneal implants displayed very minimal cellular infiltration into the material itself at 7- and 21-days post-implantation, with clusters of cellular infiltrates seen more frequently than a diffuse infiltrate as observed in SQ and VML implants (**Fig. 5a**). In SQ implants, there is an increase in cellular infiltration into the material itself observed by 21 days post-implantation. In contrast, the cellular response was primarily associated with surrounding tissue at 7 days post-implantation. In VML implants, there was cellular infiltration around damaged muscle fibers which continued into the scaffold area with no strong fibrotic encapsulation or border around the material. When counting the cellular infiltration by collagenase digestion and isolation of single cells from the material and surrounding tissue space, we found that the injury induced the highest cellular infiltration into the materials and tissue, followed by intraperitoneal implants (**Fig. 5b**). When instead counting by histological images and focusing on the material area, we could see a much stronger infiltration into the material itself in VML implants compared to IP and SQ (3491 cells/mm^2^ VML, 548 cells/mm^2^ IP, 427 cells/mm^2^ SQ), with the SQ increasing by 21 days post-implantation (**Fig. 5c;** 368 versus 1439 cells/mm^2^). When evaluating the IP implant, there is a relative decrease in cellular infiltration when the surrounding tissue is excluded from analysis, as the flow cytometric evaluation includes a lavage of the peritoneal cavity to encompass cells within the tissue space in response to the material implantation.

**FIG 5.**
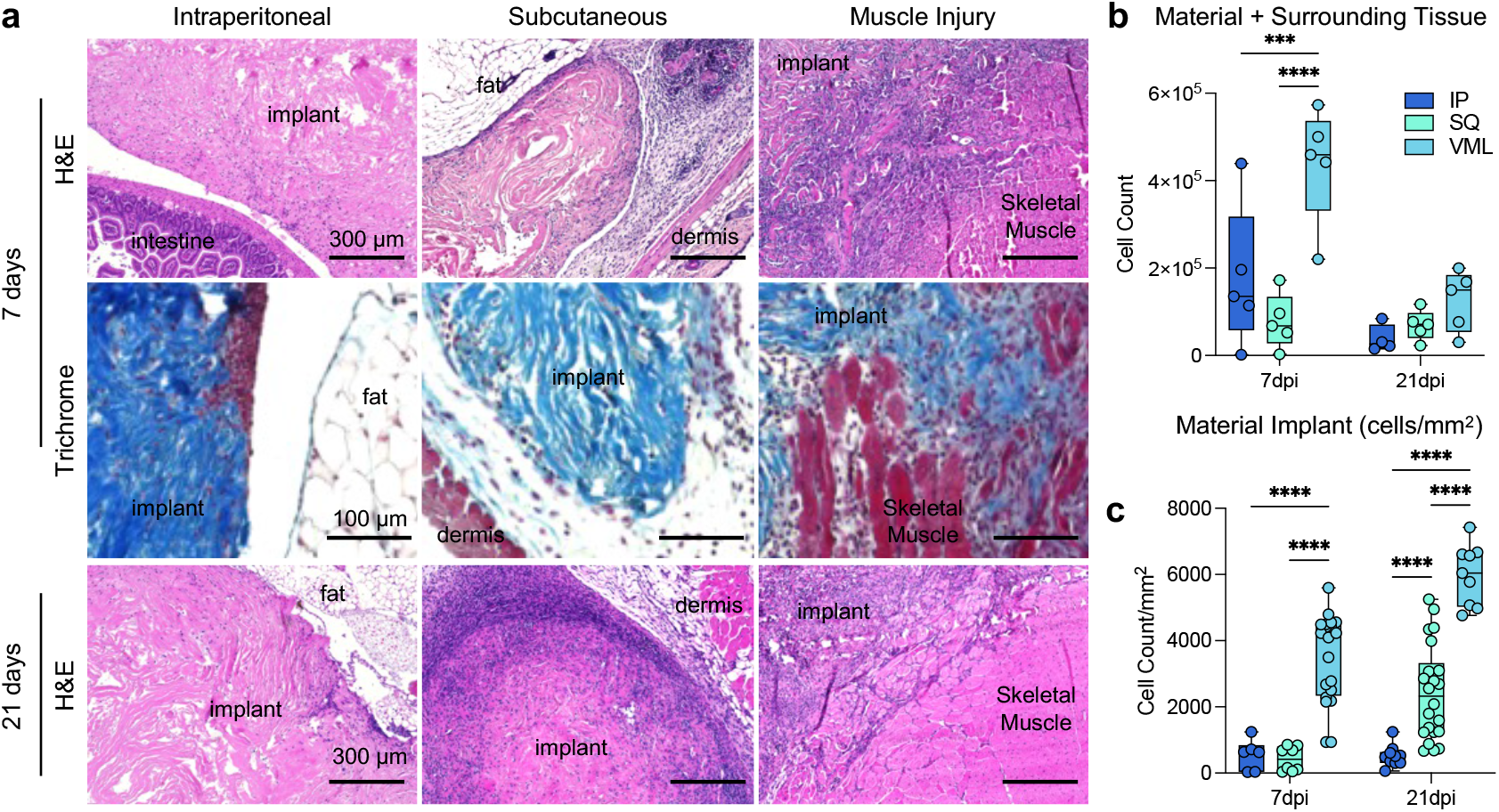
Histopathologic differences in cellular infiltration to ECM implants. (**a**) Top row: hematoxylin and eosin (H&E) staining of IP (intraperitoneal), SQ (subcutaneous) and VML (muscle injury) implants at 7 days post-implantation (dpi). Middle: Masson’s trichrome of tissue interface at 7dpi. Bottom row: H&E of IP, SQ, and VML implants at 21 dpi. Representative of n = 5 mice. (**b**) Count of cells by isolation and flow cytometry (**c**) Count of cells per mm^2^ of material only (not including surrounding tissue). Data are range, n = 5 mice, ANOVA with Tukey posthoc. *** = p < 0.001; **** = p < 0.0001.

## 5. DISCUSSION

As biomaterials are used in numerous tissue contexts, evaluating them in preclinical models that reflect this diversity of applications is important. The immune microenvironment of different tissues is dependent upon a number of factors including the unique nature of tissue-resident immune cells [16], the secretome of stromal and parenchymal cells [17], the differences in microbiome constituents [23], and the history of past infection or injury [24]. Using a head-to-head analysis of the same biomaterial placed in different tissue microenvironments, we uncovered several conserved immune response profiles to ECM scaffolds and tissue-specific characteristics in this study.

As expected, macrophage infiltration was a common response to all implanted biomaterials, regardless of tissue locations. Macrophages play an essential role in the foreign body response [25, 26]. When a biomaterial is implanted in the body, macrophages are among the first immune cells to interact with the material. Following implantation, macrophages are recruited to the site of the biomaterial through chemotactic signals released by injured tissues, immune cells, or the biomaterial itself. Macrophages can originate from circulating monocytes or resident macrophages in the surrounding tissue. In this study, we found that CX3CR1, a chemokine receptor primarily expressed on the surface of immune cells, including monocytes, macrophages, dendritic cells, etc., may play a role in cell trafficking and immune responses regarding temporal dynamics and recruitments at different locations. CX3CR1^+^ cells had opposite patterns in IP compared to SQ and VML implants, with an early preference for CX3CR1^+^ monocytes and eosinophils. This pattern was found early in SQ and VML environments and delayed in IP implants. CX3CR1 is the receptor for Fractalkine, which is an important chemotactic agent and also induces cellular adhesion; CXC3R1 has been associated with some type-2 immune responses, especially in the lung but also in the intestine and brain [27-29]. The SQ implants also recruited more Ly6C^hi^ macrophages and CD11b^+^ basophils than IP implants, with Ly6C^hi^ macrophages being associated with inflammatory peripheral blood recruitment as opposed to local tissue-resident inflammation [30, 31]. The intraperitoneal space is known to have a large abundance of tissue-resident macrophages, specifically GATA6-expressing macrophages that are the first to respond to an injury site (before neutrophils) within hours of tissue damage [32, 33]. This correlates with our findings of IP implants having a lower fraction of peripheral blood-derived macrophages when compared to areas such as the subcutaneous space.

Once macrophages have localized to the site of the ECM biomaterial, they undergo activation. Macrophage activation and polarization can result in both M1-like and M2-like macrophages, which have distinct functions and characteristics [34]. Canonically, M1-like macrophages are more anti-bacterial and inflammatory and contribute to recruiting other immune cells to the implant site. As the immune response progresses and inflammation subsides, M2-like macrophages become more predominant. M2-like macrophages contribute to tissue healing and repair by promoting angiogenesis, remodeling the ECM, and facilitating tissue integration with the biomaterial; however, it is important to note that cells rarely fall into such a binary *in vivo*, and their phenotype is tailored to both the challenge they are faced with and their tissue location [15]. As previously described in the literature, ECM materials induced an M2-like phenotype [35, 36]. This was enhanced by injury-induced, which likely compounded a type-2 response characteristic of wound healing. This was predominantly present in the expression of CD206 and CD301b. CD206 has previously been associated with positive outcomes in ECM scaffold remodeling; furthermore, it has been implicated directly in the uptake of collagen fragments in tumors suggesting a direct mechanism by which it is assisting in scaffold integration [37, 38]. CD301b plays a role in the phagocytosis of Gal-GalNAc-modified proteins for presentation to CD4^+^ T cells which have been previously shown to play a role in biomaterial-mediated muscle regeneration [5]. CD301b is also associated with positive outcomes in wound healing [39]. CD206 and CD301b were also detected on cDC1s as previously described in VML, but the co-stimulatory molecule CD86 was higher on these cells in IP implants.

In addition to macrophage recruitment, eosinophils are present in response to ECM-derived material implantations [19]. Although eosinophils are typically known to be involved in immune responses associated with allergies [40, 41], chronic inflammation [42, 43], and parasitic infections [44, 45]; the response to ECM biomaterials also involves eosinophils [7, 46]. In this study, eosinophil infiltration was stronger in the subcutaneous space when compared to the intraperitoneal space. These anatomical regions have distinct microenvironments with different cellular compositions, contributing to differences in eosinophil distribution. One possible explanation is that the subcutaneous area is more exposed to environmental allergens, which may lead to increased eosinophil infiltration compared to the intraperitoneal cavity. Furthermore, the subcutaneous space is more prone to fibrotic collagen-based healing to promote closure of the barrier tissue and prevent subsequent infections from the environment.

Regarding antigen presentation, all materials recruited a robust population of MHCII^+^ macrophages. The communication between CD4^+^ T cells and macrophages has been previously established in VML [5]. Interestingly, all ECM implants recruited a specific cDC1 population regardless of tissue location. We previously described these cells as important potential mediators in self-tolerance after injury and material implantation that are recruited downstream of damage associated molecular pattern (DAMP) engagement [19]. The fact that these cells are recruited to ECM scaffolds even in the absence of injury suggests that the material implant inherently recruits to the biomaterial microenvironment. When evaluating the data on MHCII^+^ cells, we found that IP implants were accompanied by a robust Lin^-^MHCII^+^ cell population that was virtually absent in other tissue locations. Due to the size in scatter and expression of MHCII, we determined that these were likely B cells; this was confirmed with a panel staining for lymphocytes, and we found that 70% of non-myeloid cells were B220^+^ B cells in IP implants, followed by T cells at a similar level to the VML application. Previous research by our lab and others has shown minimal B cell infiltration into ECM scaffolds in VMLs [19, 47]. This shift in dominant antigen-presenting cell types could alter downstream responses and should be considered in designing biomaterials destined for applications in the peritoneal cavity. B cells can be activated to induce antigen-presenting pathways through the engagement with extracellular antigens on the B cell receptor (BCR), which then induces upregulation of antigen processing and presentation machinery such as CD86; this process is thereby independent of other APCs such as dendritic cells [48]. B cells are generally activated by IL-4, which is necessary for their maturation and survival, and this cytokine is greatly induced by T cells in response to ECM scaffolds [5, 46]. Furthermore, B1 B cells in the peritoneal cavity have been shown to have phagocytic capabilities and help present antigens to CD4+ T cells more efficiently than macrophages [49]. B cell antigen presentation has also been described in tolerance in multiple models and has been associated with suppressor CD8+ T cells in the eye [50]; recently, we have shown that CD8+ regulatory T cells may be involved in response to injury and ECM biomaterial implantation [19].

This work highlights the divergent immune phenotypes depending on tissue location, emphasizing the need for appropriate preclinical models of biomaterial implantation. Of great interest, the robust difference in B cell infiltration in the peritoneal cavity suggests that this tissue location strongly enriches both cellular and humoral arms of adaptive immunity. This observation may have implications for downstream applications of materials or revision surgeries. As a result, the tissue context where the biomaterial is implanted and the presence or absence of an injury play an important role in the outcome of that material; hence, scaffolds should be tuned to their specific applications to promote integration and tissue regeneration.

## 7. ACKNOWLEDGEMENTS & FUNDING

The authors thank Vanathi Sundaresan for her assistance in laboratory organization. This work was funded entirely by the Intramural Research Program of the National Institute of Biomedical Imaging and Bioengineering within the National Institutes of Health. Author contributions: SD, AJ, MF, and DF completed experiments; RL, TBN, and KS analyzed data. TBN and KS wrote the manuscript. <u>Disclaimer</u>: The content of this publication does not necessarily reflect the views or policies of the Department of Health and Human Services, nor does mention of trade names, commercial products, or organizations imply endorsement by the U.S. government. The NIH, its officers, and employees do not recommend or endorse any company, product, or service.

## 8. DATA AVAILABILITY

All data in this manuscript are available upon request from the authors.

