## Supplemental Materials for "Conserved and tissue-specific immune responses to biologic scaffold implantation"

### SUPPLEMENTARY FIGURES AND LEGENDS

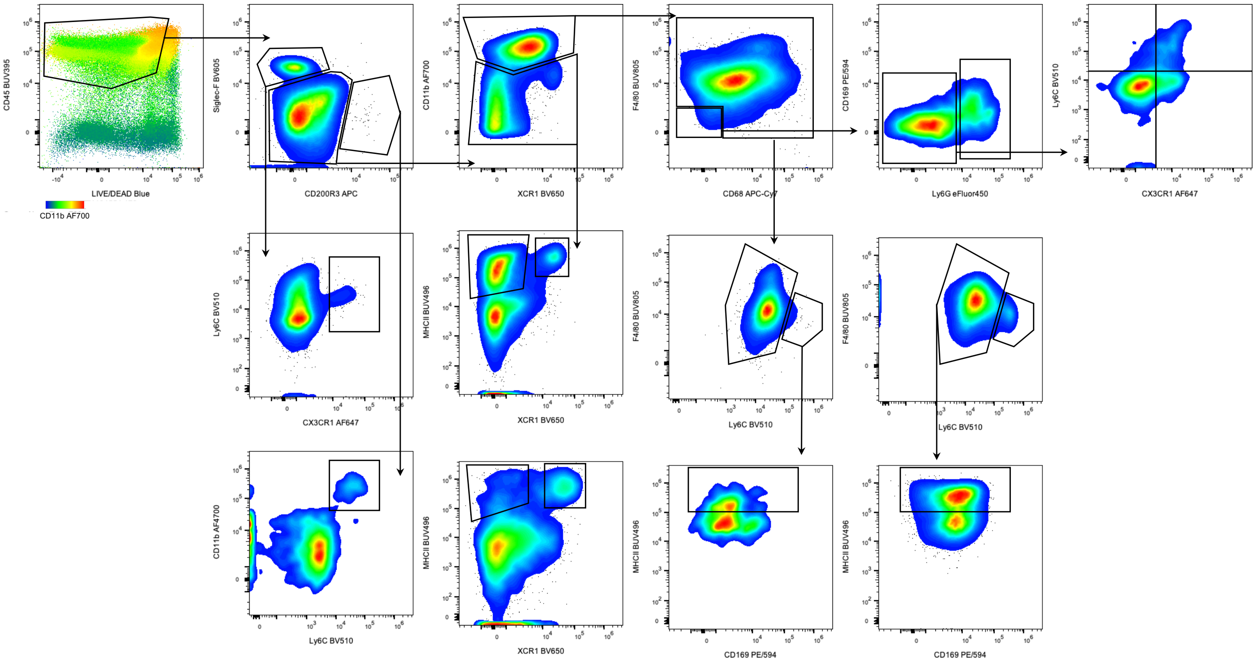


**SUPPLEMENTARY FIGURE 1 | Flow cytometry gating strategy for myeloid phenotyping.** Example gating is shown from intraperitoneal and volumetric muscle loss implant samples. FSC-A x SSC-A and FSC-A x FSC-H were used to discriminate debris and doublets before gating on live immune cells (FSC = forward scatter, SSC = side scatter, A = area, H = height). Modified from a preprinted manuscript [19].

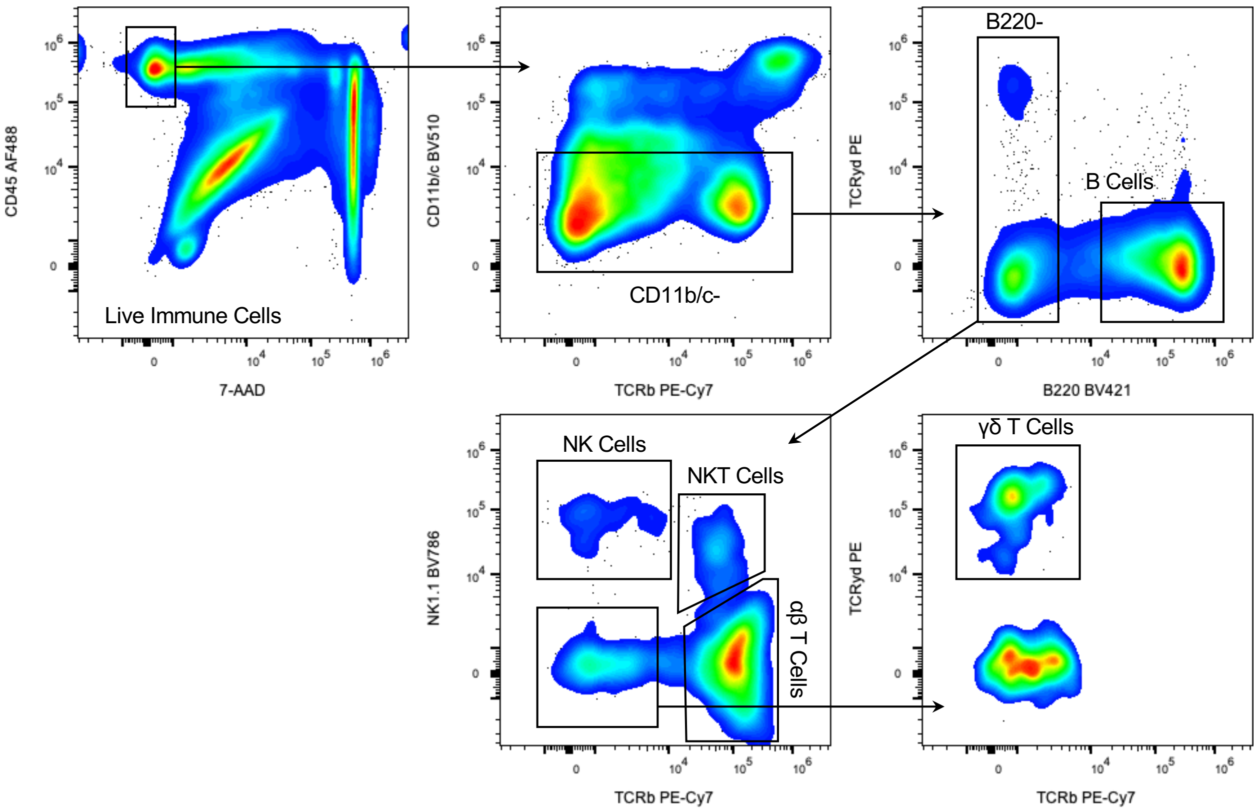


**SUPPLEMENTARY FIGURE 2 | Flow cytometry gating strategy for lymphoid phenotyping.** Example gating shown from intraperitoneal implant sample. FSC-A x SSC-A and FSC-A x FSC-H were used to discriminate debris and doublets before gating on live immune cells (FSC = forward scatter, SSC = side scatter, A = area, H = height).

| Marker | Target | Clone | Fluorophore | Vendor | Dilution Factor | Bead or Cell Control |
|---|---|---|---|---|---|---|
| CD45 | Leukocytes | 30-F11 | BUV395 | BD | 1:100 | PBMC |
| Viability | Dead Cells | | LIVE DEAD Blue | Thermo | 1:1000 | RAW264.7 |
| MHCII | APC | 2G9 | BUV496 | BD | 1:400 | PBMC |
| CD1d | Lipid Ag Present | 1B1 | BUV563 | BD | 1:200 | Beads |
| F4/80 | Macrophage | T45-2342 | BUV805 | BD | 1:100 | Beads |
| CD206 | M2 Mac | C068C2 | BV421 | Biolegend | 1:100 | Beads |
| Ly6G | Neutrophil | 1A8 | eFluor 450 | Thermo | 1:200 | Beads |
| CD103 | DC | M290 | BV480 | BD | 1:200 | Beads |
| Ly6C | Monocyte | HK1.4 | BV510 | Biolegend | 1:200 | PBMC |
| CD11c | DC | N418 | BV570 | Biolegend | 1:200 | Beads |
| Siglec-F | Eosinophil | E50-2440 | BV605 | BD | 1:200 | Beads |
| XCR1 | DC | ZET | BV650 | Biolegend | 1:400 | Beads |
| CD11b | Myeloid | M1/70 | Alexa Fluor 700 | Biolegend | 1:400 | PBMC |
| CD8α | DC | 53-6.7 | Spark Blue 550 | Biolegend | 1:100 | PBMC |
| CD301b | Macrophage | URA-1 | PE/Cy7 | Biolegend | 1:400 | PBMC |
| CD86 | M1 Mac | A17199A | PE | Biolegend | 1:400 | PBMC |
| CD169 | Tissue Resident | 3D6.112 | PE/Dazzle594 | Biolegend | 1:400 | Beads |
| CCR7 | M1 Mac | 4B12 | PE-Cy5 | Biolegend | 1:200 | PBMC |
| CD200R3 | Baso/Mast Cell | Ba13 | APC | Biolegend | 1:200 | Beads |
| CX3CR1 | Pre-M2 | SA011F11 | Alexa Fluor 647 | Biolegend | 1:400 | PBMC |
| CD68 | Macrophage | FA-11 | APC-Cy7 | Biolegend | 1:100 | Beads |

**SUPPLEMENTARY TABLE 1 | Flow cytometry antibodies for the myeloid phenotyping panel.** Modified from a preprinted manuscript [19].

| Marker | Target | Clone | Fluorophore | Vendor | Dilution Factor | Bead or Cell Control |
|---|---|---|---|---|---|---|
| CD45 | Immune | I3/2.3 | AF488 | BD | 1:100 | Beads |
| Viability | Dead Cells | | 7-AAD | BD | Neat | PBMCs |
| TCR γδ | γδ T Cells | GL-3 | PE | BD | 1:100 | Beads |
| TCRβ | αβ T cells | H57-597 | PE/Cy7 | BD | 1:100 | Beads |
| B220 | B Cells | RA3-6B2 | BV421 | BD | 1:100 | Beads |
| CD11b | Myeloid | M1/70 | BV510 | BD | 1:100 | Beads |
| CD11c | Myeloid | HL3 | BV510 | BD | 1:100 | Beads |
| NK1.1 | NK Cells | PK136 | BV786 | BD | 1:100 | Beads |

**SUPPLEMENTARY TABLE 2 | Flow cytometry antibodies for a lymphoid phenotyping panel.** Modified from a preprinted manuscript [19]
